# 24-nt phasiRNAs move from tapetal to meiotic cells in maize anthers

**DOI:** 10.1101/2021.08.24.457464

**Authors:** Xue Zhou, Kun Huang, Chong Teng, Ahmed Abdelgawad, Mona Batish, Blake C. Meyers, Virginia Walbot

## Abstract

- In maize, 24-nt phased, secondary small interfering RNAs (phasiRNAs) are abundant in meiotic stage anthers, but their distribution and functions are not precisely known.
- Using laser capture microdissection, we analyzed tapetal cells, meiocytes, and other somatic cells at several stages of anther development to establish the timing of 24-*PHAS* precursor transcripts and the 24-nt phasiRNA products.
- By integrating RNA and small RNA profiling plus single-molecule and small RNA FISH (smFISH or sRNA-FISH) spatial detection, we demonstrate that the tapetum is the primary site of 24-*PHAS* precursor and *Dcl5* transcripts and the resulting 24-nt phasiRNAs. Interestingly, 24-nt phasiRNAs accumulate in all cell types, with the highest levels in meiocytes, followed by tapetum.
- Our data support the conclusion that 24-nt phasiRNAs are mobile from tapetum to meiocytes and to other somatic cells. We discuss possible roles for 24-nt phasiRNAs in anther cell types.

## Introduction

In plants, small RNAs (sRNAs) belong to three main categories: microRNAs (miRNAs) involved in target mRNA cleavage, heterochromatic small interfering RNAs (hc-siRNAs) involved in RNA-directed DNA methylation (RdDM), and phased, secondary small interfering RNAs (phasiRNAs). PhasiRNAs were initially reported in male reproductive organs of maize and rice (Johnson *et al*., 2009; Song *et al*., 2012a; Zhai *et al*., 2015; Fei *et al*., 2016; Tamim *et al*., 2018; Teng *et al*., 2020), and more recently in diverse monocots and eudicots (Kakrana *et al*., 2018; Xia *et al*., 2019; Patel *et al*., 2021; Pokhrel *et al*., 2021); they are found primarily in anthers. In most species, 21-nt phasiRNAs predominate during early anther development, while 24-nt phasiRNAs are abundant during meiosis.

PhasiRNA biogenesis starts with transcription of long noncoding precursors (*PHAS* transcripts) by RNA polymerase II. Subsequently, the *PHAS* transcripts are capped and polyadenylated, translocated to the cytoplasm, and then bound by ribosomes (Yang *et al*., 2021). In the polysomal complex, an Argonaute-associated 22-nt miRNA complementary to sequences in the *PHAS* transcript binds to this precursor molecule and triggers cleavage. miR2118 is utilized for most 21-nt *PHAS* precursors and miR2275 for most 24-nt *PHAS* precursors based on the presence of complementary sequences; a fraction of precursors lacks such complementarity and may be processed utilizing other miRNAs. Then, the 3’ portions of cleaved transcripts are converted to double-stranded RNA by RNA-DEPENDENT RNA POLYMERASE 6 (RDR6), and subsequently processed by Dicer-like proteins (DCL4 for the 21-nt and DCL5 for the 24-nt type) to yield the phasiRNAs (Song *et al*., 2012a; Song *et al*., 2012b; Zhai *et al*., 2015; Teng *et al*., 2020). Despite the expectation of equal stoichiometry of all the products from one *PHAS* transcript, individual phasiRNA abundances derived from the same transcript can differ by more than 1000-fold indicating differential stabilization (Tamim *et al*., 2018; Zhang *et al*., 2021). It is possible that only these few abundant products generated from one precursor have functions.

In maize the 21-nt phasiRNAs are distributed throughout immature anther lobes based on *in situ* hybridizations with one or a few examples, although the production of miR2118 is restricted to the epidermis (Zhai *et al*., 2015). Several observations associate phasiRNAs with male sterility. First, the absence of specific 21-nt phasiRNAs are implicated in causing male sterility in rice (Komiya *et al*., 2014; Fan *et al*., 2016). Second, in numerous male-sterile mutants of maize, either the 21-nt or 24-nt phasiRNAs are absent reflecting the tight control of the normal timing of their appearance during development (Zhai *et al*., 2015; Nan *et al*., 2017; Teng *et al*., 2020; Yadava *et al*., 2021). More specifically addressing the role of 24-nt phasiRNAs, *dicer-like5* mutants unable to process most 24-nt *PHAS* precursors are male-sterile and exhibit aberrant tapetal development under normal growing conditions (Teng *et al*., 2020). In maize, the expression of *PHAS* precursors and miR2275 as well as the accumulation of 24-nt phasiRNAs occur mainly in the tapetum at the prophase I stage in anthers (Zhai *et al*., 2015). Male-sterile mutants with tapetal developmental defects evident during early meiotic stages are missing the 24-nt phasiRNAs (Nan *et al*., 2017; Teng *et al*., 2020). Interestingly, the male sterility cases in rice and maize exhibit temperature-dependent male sterility: in appropriate growing conditions, the plants are male-fertile despite the absence of specific or an entire class of phasiRNAs (Fan *et al*., 2016; Teng *et al*., 2020; Yadava *et al*., 2021). This observation has led to the hypothesis that phasiRNAs may buffer anther development against environmental perturbation, particularly in the tapetum. In addition to the expectation that phasiRNAs could modulate mRNA half-life of still poorly defined targets, the 24-nt phasiRNAs result in increased CHH DNA methylation at 24-*PHAS* loci in maize, as assessed in whole anthers (Zhang *et al*., 2021). In analysis of isolated meiocytes, 24-nt phasiRNAs have been detected, and there is an increase in CHH DNA methylation, leading to the hypothesis that these phasiRNAs might regulate meiosis (Dukowic-Schulze *et al*., 2016) by an unknown mechanism related to *cis* methylation of their own loci. To date, the precise role of 24-nt phasiRNAs in regulating maize anther development has remained elusive.

sRNAs can function by travelling within and between organs: from the site of synthesis, they can spread both to neighboring cells and systemically over long distance (Chitwood & Timmermans, 2010; Dunoyer *et al*., 2010; Molnar *et al*., 2010). Cell-to-cell movement of sRNAs is probably through plasmodesmata (PD), a plasma membrane-lined pore acting as an intercellular channel that connects the cytoplasm of adjacent cells (Dunoyer *et al*., 2013). To date, the potential movement of 24-nt phasiRNAs from the site of biogenesis to the neighboring cell layers in maize anthers has not been well studied. A major concern with our previous localization analysis for 24-nt phasiRNAs and biogenesis components primarily to the tapetum is that only a few examples were evaluated, and the probes could adhere to the callose coat surrounding the germinal cells, yielding a non-specific signal (Zhai *et al*., 2015). Therefore, in meiotic stage maize anthers, two major questions should be addressed: 1) do meiocytes contain all the components for generating 24-nt phasiRNAs? And 2) using RNA-seq and small RNA-seq, what is the timing of synthesis and the distribution of these components in anther lobes?

To explore the distribution of 24-nt phasiRNA biogenesis components and the resulting small RNAs, we used laser capture microdissection (LCM) to isolate meiocytes (ME), tapetum (TAP), and other somatic cells (OSC) from two anther developmental stages for low input RNA-seq and sRNA-seq and for a more comprehensive developmental analysis using RT-qPCR. By integrating comprehensive RNA/sRNA profiling and single-molecule or sRNA FISH (smFISH/sRNA-FISH) spatial detection, we demonstrate that TAP contain the most 24-*PHAS* primary transcripts, whereas ME ultimately accumulate more 24-nt phasiRNAs at meiotic prophase I. We discuss these results in a model in which 24-nt phasiRNAs move from TAP to ME during meiosis.

## Materials and Methods

### Plant material

Maize (*Zea mays)* W23 *bz2* inbred line plants were grown in Stanford, CA, USA under greenhouse conditions (31 °C/21 °C, 14 h day/10 h night light cycle with supplemental LED and UV-A lamps providing ~50% of summer sun conditions). Anthers were dissected from tassels and then measured using a micrometer (Fisher Scientific). Homozygous *dcl5-mu03//dcl5-mu03* plants are temperature-sensitive male sterile. They are male sterile under optimized growth condition (28 °C/22 °C day/night temperature, 14 h light, 400 μmol/m^2^/s) in the greenhouse. *dcl5-mu03//dcl5-mu03* male sterile and *dcl5-mu03//Dcl5* fertile siblings were grown in St. Louis, MO, USA under the optimized growth conditions in a greenhouse.

### Laser Capture Microdissection (LCM)

Fifty upper floret anthers were dissected from individual central tassel spikes for each biological replicate. Different stage anther samples were collected from different maize plants. ME, TAP, and OSC were isolated from 1.25 mm, 1.5 mm, and 2.0 mm W23 anthers by LCM as described previously (Kelliher & Walbot, 2012), with minor modification. Fixed anthers were cryoprotected in 10% (2-3 days) and 15% sucrose/PBS (phosphate buffered saline) (5 to 7 days) to better preserve tapetal cell layer morphology. Total RNA was isolated from each cell type using the RNAqueous™-Micro Kit (Invitrogen); RNA quality was evaluated on an Agilent Bioanalyzer using the RNA 6000 Pico Kit (Agilent Technologies).

### Cytological staging and imaging

AR/PMC/ME from W23 anthers (from the 0.75 mm through 2.25 mm anther length stages, recovered at 0.25 mm intervals) were extruded, classified using a micrometer, and then appropriately sized anthers were stained with 10 μg/mL Hoechst 33342 (Sigma-Aldrich) according to our pervious report (Nelms & Walbot, 2019). Confocal image stacks for the germinal cells were taken with a Leica SP8 microscope using a 93X Glycerol immersion objective. ImageJ (Schneider *et al*., 2012) was used for processing the fluorescent microscopy images. Cytological stages during meiotic prophase I were assigned using established criteria (Dawe *et al*., 1994; Nelms & Walbot, 2019).

### RNA-seq and sRNA-seq library construction and sequencing

Total RNA was isolated from three different LCM cell samples and FA samples at 1.5 mm and 2.0 mm using the RNAqueous™-Micro Kit (Invitrogen). RNA-seq libraries were prepared using CEL-seq2 (Hashimshony *et al*., 2016) based on a modified protocol (Nelms & Walbot, 2019). Primers were synthesized by phosphoramidite chemistry at the Stanford Protein and Nucleic Acid Facility (Stanford, CA) (Table **S1**). All RNA-seq libraries were sequenced on an Illumina HiSeq 4000 instrument at the Stanford Genome Sequencing Service Center (Stanford, CA) with paired-end 150 bp reads.

For small RNA library preparation, RealSeq®-AC kits (SomaGenics) were used for ultra-low amounts of total RNA starting with 100 ng from whole anthers or 10 ng from LCM cell samples. Sequencing in a single-end mode on an Illumina NextSeq (University of Delaware) yielded 75 bp reads for sRNA-seq.

### Data handling and bioinformatics

Paired-end RNA-seq Read1 contained a 10 bp unique molecular identifier (UMI) (Kivioja *et al*., 2011) and sample barcodes, and Read2 contained the transcript sequence. Each UMI was processed and attached to the Read2 metadata using fastp (Chen *et al*., 2018). Barcodes were used to demultiplex Read2 into a separate file for each library using fastq-multx (Aronesty, 2013). After trimming the adapters by Trim Galore, the reads were mapped to version 4 of the B73 maize genome using Hisat2 (Pertea *et al*., 2016). RNA-seq reads were normalized by dividing by the total number of transcripts (UMIs) in each sample and multiplying by one million (transcripts per million normalization).

sRNA-seq data were processed as previously described (Mathioni *et al*., 2017). In brief, Trimmomatic version 0.32 (Bolger *et al*., 2014) was used to remove the linker adaptor sequences. Then, the trimmed reads were mapped to version 4 of the B73 maize genome using Bowtie (Kersey *et al*., 2016). Read counts were normalized to one million to allow for the direct comparison across libraries. PhasiRNAs were designated based on the criteria of a 24-nt length and mapping coordinates within the previously identified 176 24-*PHAS* loci (Zhai *et al*., 2015) that were updated to version 4 of the B73 genome using the assembly converter tool (Tello-Ruiz *et al*., 2018).

### Quantitative reverse transcription PCR (RT-qPCR)

RT-qPCR for multiple stage anthers (0.75 mm – 2.0 mm; 0.25 mm intervals) and LCM cell sample collections (ME, TAP, and OSC) was performed using the Luna® Universal Probe One-Step RT-qPCR Kit (New England Biolabs). Quantitative PCR was performed using TaqMan primers synthesized by Integrated DNA Technologies (Table **S2**) on a CFX96 C1000 Touch Real-Time PCR Detection System (BioRad). *ZmCyanase* expression was used to normalize among biological samples, because this gene is highly expressed at all stages of anther development (Fig. **S5**). Each sample type was tested in two biological and three technical replicates.

### Single-molecule fluorescence *in situ* hybridization (smFISH) for mRNAs and sRNAs

Maize spikelets of various sizes were dissected from the central spike and fixed in 20 mL glass vials using 4% paraformaldehyde in 1x PHEM buffer (5 mM HEPES, 60 mM PIPES, 10 mM EGTA, 2 mM MgCl2 pH 7). Fixation was done three times in a vacuum chamber at 0.08 MPa, 15 min each. After fixation, samples were sent for paraffin embedding at the histology lab in the Nemours/Alfred I. duPont Hospital for Children (Wilmington, DE). Embedded paraffin samples were sectioned using a paraffin microtome and dried on poly-l-lysine-coated 22 × 22 mm #1.5 coverslips (Carl Zeiss Microscopy, LLC, Cat# 474030-9020-000). Samples were then de-paraffinized using Histo-Clear (Fisher Scientific, 50-899-90147) and re-hydrated by going through an ethanol series of 100, 95, 80, 70, 50, 30, 10% (vol/vol) (30 sec each) and finally water (1 min) at room temperature. After protease (Sigma, P5147) digestion (20 min, 37°C), samples were neutralized in 0.2% glycine (Sigma-Aldrich, G8898) for 2 min. After two washes in 1X PBS buffer, samples were dehydrated and then hybridized with smFISH probes. About 20-nt smFISH probes were designed to bind specifically across the length of each target RNA in a non-overlapping series (Markey *et al*., 2014). For small RNA detection, probes were designed reverse complemented to ~19 nt of the phasiRNA sequences generated from each locus. Briefly, 23-51 probes (total coverage of about 800 nt for mRNAs or sRNAs) were designed for each target, and each probe was synthesized with a 3’ amino modification (LGC Biosearch Technologies, CA) (Table **S3**). All the probes of one set were pooled and *en masse* coupled with Alexa Fluor 647 (Thermo Fisher) or Texas Red; the labeled probe fraction was purified using reverse phase HPLC (Batish *et al*., 2011). The probes were diluted to a concentration of 20 ng/μl in a hybridization buffer containing 10% formamide, 1 mg/mL yeast tRNA, 10% dextran sulfate, and 2 mM Vanadyl ribonucleoside complex (New England Biolab, Ipswich, MA, Catalogue Number, S142), 0.02% RNase free BSA (Ambion, AM2618). The biological samples were hybridized with probes overnight in a humid chamber at 37°C. The samples were then washed two times with 2X SSC buffer (sodium saline citrate) containing 10% formamide, and a final wash was done using 1X TBS buffer (Tris-buffered saline). Samples were mounted using ProLong™ Glass Antifade Mountant with NucBlue™ Stain (ThermoFisher Scientific, P36981).

Fluorescence was detected by spectral unmixing of autofluorescence spectra using laser scanning confocal microscopy on a Zeiss LSM 880 multiphoton confocal microscope. We used 30% power of the 633 nm laser and an alpha Plan-Apochromat 100x/1.46 oil lens to acquire RNA *in situ* images. Pure dye was used as positive control, and autofluorescence from maize tissue was used as the negative control for spectral bleed-through. After images were acquired, each image was spectra unmixed. The brightness and contrast of images in the same figure panel were adjusted equally and linearly in Zen 2010 (Carl Zeiss). For quantification, the numbers for each localization event were calculated using ImageJ (SpotQuant). Two replicates for each Z-stack of images were used for calculating the copy number for each gene’s transcripts.

## Results

### LCM collections from meiotic stage maize anthers

Control of fertility in maize anthers, the male reproductive floral organs, is important for hybrid seed production as an economic way to improve grain yield. Maize tassels produce hundreds of spikelets, each of which contains an upper and lower floret, and each floret contains three anthers (Fig. **1a**). The lower floret stamens develop approximately 1 day later than the upper floret stamens within the same spikelet. At the beginning of meiosis, each anther contains four lobes; each lobe consists of four somatic cell types that are required to support meiotic cell development (Fig. **1a**). The high regularity of maize anther development, numerous anthers, and a detailed staging system allow rapid dissection of carefully staged anthers for cytological and molecular analysis (Fig. **1b, c**) (Kelliher & Walbot, 2011). To explore the spatial distribution of 24-nt phasiRNAs and their pathway components in the W23 inbred (Fig. **2a**), LCM was used to isolate three samples (ME, TAP, and OSC) from two stages of meiotic anthers (1.5 mm zygotene and 2.0 mm pachytene) (Fig. **S1**; Table **S4**). Collections from additional stages were used in RT-qPCR analysis to provide finer detail of the timing of biogenesis of a subset of 24-*PHAS* precursor and *Dcl5* transcripts.

**Fig. 1.**
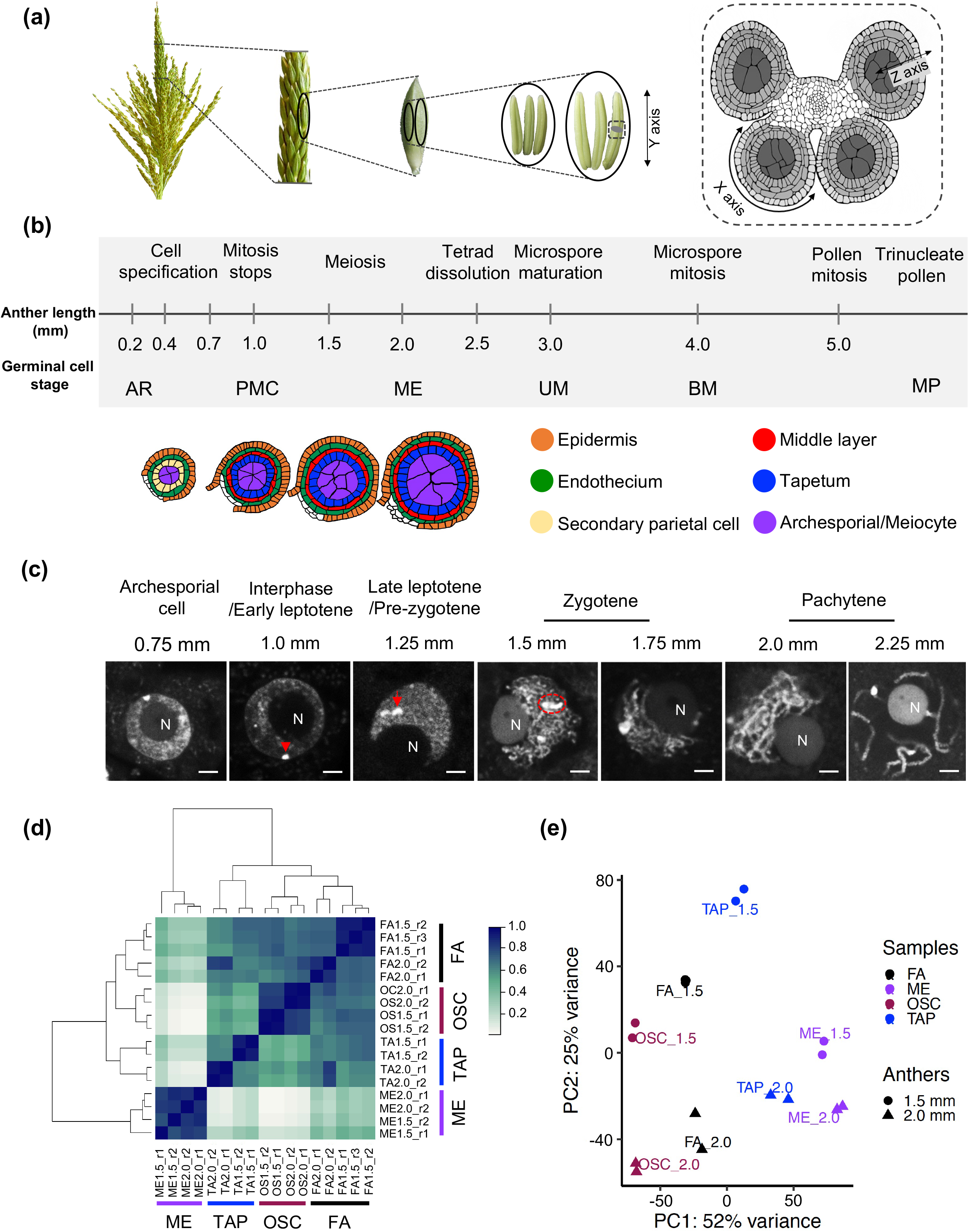
Staging of maize anthers and quality evaluation of LCM cell sample collections. (**a**) Male inflorescence structures. From left to right: maize tassel; central spike; single spikelet; three lower and three larger upper florets; transverse confocal microscopy view of a single anther (near the start of meiosis). (**b**) The timeline of maize anther development from initial cell specification, proliferation, and differentiation to pollen maturation. Approximately 10 days of development occur between anther initiation and the start of meiosis at 1.2 mm; meiosis requires 6 days and finishes at 2.5 mm with the tetrad stage; an additional 15-16 days are required for microspore and then trinucleate pollen development (note that this portion of the timeline is not drawn to scale), followed by pollen maturation and anthesis after 5 more days. Tracings of confocal images of single anther lobes are shown through the dyad stage. The germinal cell stages are abbreviated as follows: mitotic archesporial cells (AR); post-mitotic pollen mother cells (PMC); meiocytes (ME); uninucleate microspore (UM); binucleate microspore (BM); and mature pollen (MP). (**c**) Fluorescence microscopy images of germinal cells visualized with Hoechst staining. During the meiotic proliferation period (0.75 mm anther), archesporial cells have a central nucleolus (N) and a smaller volume than meiotic nuclei. At premeiotic interphase/ early leptotene, the knobs (arrowhead) are round, the nucleoli are located centrally, and chromatin threads are not visible or visible only as localized patches. During leptotene leading to pre-zygotene, the spherical knobs become elongated (arrowhead) coincident with nucleolar movement to the periphery. During zygotene, the knobs are paired (red dashed oval); chromosomes are observed as condensation proceeds. Fully paired bivalents (4 strands per thread) and finally a centrally located nucleolus were observed at pachytene. Scale bar represents 2 μm. (**d**) Sample-sample correlation analysis in fixed anthers and LCM cell sample collections at the 1.5 mm and 2.0 mm stages. As shown in the heatmap, the biological replicates of the same sample type are highly correlated at both stages. The scale based on the Pearson correlation is from lightest (no correlation) to darkest (1.0 correlation). Abbreviations: fixed anthers (FA); meiocytes (ME); tapetum (TAP); and other somatic cells (OSC). (**e**) Principal component analysis (PCA) to evaluate the similarity of biological replicates.

**Fig. 2.**
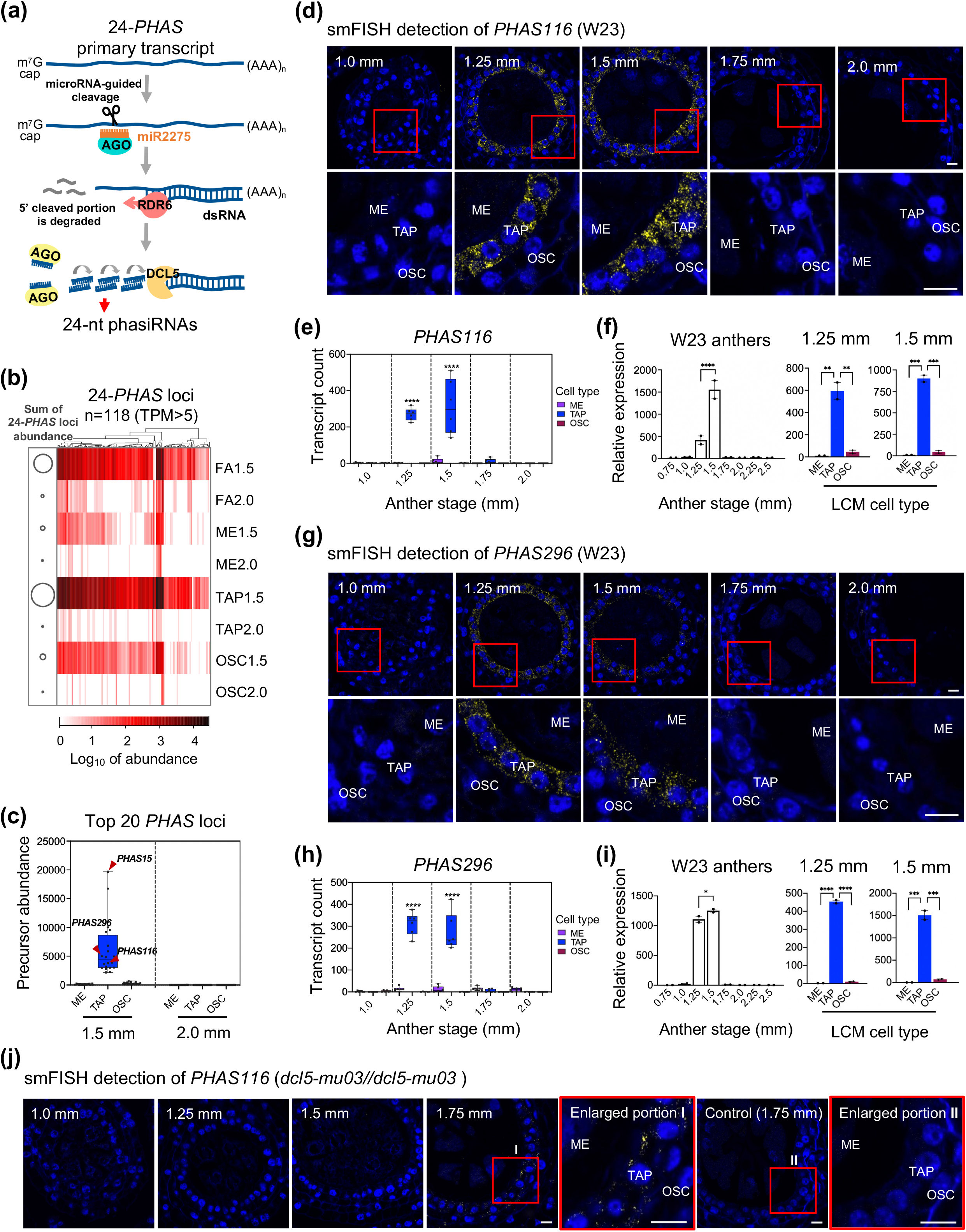
24-*PHAS* precursor distribution in maize anthers. (**a**) 24-nt phasiRNA biogenesis pathway. miR2275 is complementary to a conserved motif in precursors, resulting in precise cleavage; cleaved products are converted to double stranded molecules by RNA-dependent RNA polymerase 6 (RDR6). Unknown Argonaute (AGO) proteins participate in biogenesis and bind at least a subset of the 24-nt phasiRNA products. (**b**) Heat map depicting the abundances of 118 24-*PHAS* precursors (TPM > 5 in at least one sample) in fixed anthers and LCM cells. Solid bubbles on the left represent the total abundance of 24-*PHAS* precursors in that sample. (**c**) Boxplot depicts the distribution of the 20 most abundant 24-*PHAS* precursors in LCM cell collections. Arrowheads point to the three 24-*PHAS* loci (*PHAS15*, *PHAS116*, and *PHAS296*) chosen to represent different abundances for RT-qPCR and single-molecule FISH (smFISH) verifications. (**d**) and (**g**) smFISH was performed using probes conjugated with dye Alexa Fluor 647 (AF647) against *PHAS116* and *PHAS296* in W23 maize anthers (1.0-2.0 mm). The fluorescent micrographs were taken using laser scanning confocal microscopy. Below the images are enlargements of area in red rectangles. Yellow smFISH spots correspond to precursor transcripts, which primarily accumulate in TAP. Scale bars = 10 μm. (**e**) and (**h**) Quantification analysis of images in (**d**) and (**g**). Transcript counts show the binding events for each cell type at different anther stages using spot count in ImageJ. Two biological and three technical replicates were performed for each stage. Statistical analysis was performed with Prism 8 software. *P* values were determined using one-way ANOVA and Tukey tests. Differences among three cell sample collections within the same anther stage were considered significant for *P* values ≤ 0.05 (* *P* ≤ 0.05; ** *P* ≤ 0.01; *** *P* ≤ 0.001; **** *P* ≤ 0.0001). (**f**) and (**I**) Quantification of localization of *PHAS116* and *PHAS296* in whole anthers (0.75- 2.5 mm) and LCM cells (1.25 and 1.5 mm) by RT-qPCR. The expression patterns of these 24-*PHAS* precursors are very similar, with high expression levels in tapetum at 1.25 mm, with a peak at the 1.5 mm stage. The tapetal cell sample is significantly higher than either meiocytes or other somatic cells. *P* values were determined using one-way ANOVA and Tukey tests in Prism 8 software. Differences among the samples were considered significant for *P* values ≤ 0.05 (* *P* ≤ 0.05; ** *P* ≤ 0.01; *** *P* ≤ 0.001; **** *P* ≤ 0.0001). Note that there is a different scale for each locus. The reference gene (*ZmCyanase)* was used to normalize expression for the accurate assessment of target 24-*PHAS* locus expression. (**j**) smFISH detection of *PHAS116* in homozygous *dcl5-mu03//dcl5-mu03* anthers. The red boxes labeled I and II are enlarged in the adjacent image to the right. No signal was detected in control hybridization assays (*Dcl5* probes) at 1.75 mm stage anthers. Scale bars = 10 μm.

### 24-*PHAS* precursors accumulate in tapetal cells

Seventeen RNA-seq libraries were built from ME, TAP, and OSC recovered from 1.5 mm and 2.0 mm stage anthers plus single, fixed anthers (FA) (Table **S5**). To check the reproducibility of the datasets, a correlation heatmap and a Principal Component Analysis (PCA) plot were constructed (Fig. **1d, e**). Biological replicates are highly similar and distinct from other cell collections at both stages, confirming the high quality of our LCM sample collections. 126 (TPM > 0) of 176 previously discovered 24-*PHAS* precursor types were identified in FA and LCM cell sample collections (Table **S6**) (Zhai *et al*., 2015); *PHAS* precursors that overlapped protein coding genes were excluded. As shown in Fig. **2b**, 118 of these loci had sufficient and reliable expression (TPM reads > 5 in at least one sample) for further analysis. Overall, the 24-*PHAS* precursors were most abundant in TAP at the 1.5 mm stage, showing an expression pattern and high abundance similar to that in 1.5 mm FA (Fig. **2b**); therefore, the majority of 24-*PHAS* precursors in whole anthers are found in the tapetal cells. The average total expression levels of 24-*PHAS* loci are approximately 65 times lower in ME and 20 times lower in OSC at this zygotene stage (Fig. **2b**; Table **S6**). Single cell RNA-seq (scRNA-seq) analysis of individual ME detected a low abundance of transcripts from 24-*PHAS* loci during the pre-zygotene/zygotene stages in a A188/B73 maize hybrid stock (Nelms & Walbot, 2022), confirmation of low precursor abundance in meiocytes. The total average abundance of these 24-*PHAS* transcripts per zygotene sample in A188/B73 is less than that in the pooled meiocyte LCM samples at 1.5 mm from W23 maize anthers. Furthermore, our previous scRNA-seq data also identified extremely low abundances of 24-*PHAS* precursors in premeiotic AR and PMC in W23 maize anthers (Nelms & Walbot, 2019). These data suggests that the TAP is the main source for 24-*PHAS* precursors in zygotene stage maize anthers.

To further verify our LCM RNA-seq data and to quantify transcripts in more stages, we randomly selected three highly expressed 24-*PHAS* precursors (*PHAS15, PHAS116*, and *PHAS296)* for smFISH and RT-qPCR assays (Fig. **2c**). In the LCM RNA-seq data, transcripts for all three loci were significantly enriched in TAP at the 1.5 mm stage compared to ME and OSC (Fig. **S2**). smFISH images confirmed that TAP contained most of these three 24-*PHAS* transcripts (yellow dots resulting from dye AF647 labeling) at the 1.25 mm and 1.5 mm stages during late leptotene/pre-zygotene and zygotene stages in W23 anthers (Fig. **2d, g**, Fig. **S3a**). Quantification counts (transcript count) demonstrated that TAP is significantly higher at both stages (Fig. **2e, h**, Fig. **S3b**). Almost no expression was detected at earlier or later stages (Fig. **2d, e, g, h**, Fig. **S3a**) or in control hybridization assays using probes that should only detect the long precursor transcript (Fig. **S4**). To confirm this assumption, hybridization assays were repeated now including seven probes that overlap sufficiently with product 24-nt small RNAs to detect phasiRNAs from *PHAS15*. The signals for *PHAS15* were highly increased when these seven overlapping probes were included; furthermore, signal was detected in three cell types including expression at the 1.75 mm stage as expected for 24-nt phasiRNAs (Fig. **S3d**). These data further verified that the distinction between 24-nt phasiRNAs and precursors can be discriminated by smFISH using our methods. For RT-qPCR using W23 anthers and LCM samples, *ZmCyanase* was used as the internal reference; it is consistently expressed across all anther stages and is expressed in all cell layers based on smFISH detection (Fig. **S5**) compared to the no label control hybridization test (Fig. **S6**). In conclusion, in 0.75-2.5 mm W23 maize anthers, the expression patterns of the three 24-*PHAS* loci showed a similar pattern, with high expression levels in both 1.25 mm and 1.5 mm anthers and with significant enrichment in TAP compared to ME and OSC (Fig. **2f, i**, Fig. **S3c**).

To further solidify the conclusion that TAP is the primary site of 24-*PHAS* transcript biogenesis, we utilized the *dcl5-mu3* mutant. Several *dcl5* mutants have been documented to contain normal 24-*PHAS* precursor levels but lack virtually all the 24-nt phasiRNAs (Teng *et al*., 2020). We reasoned that in the absence of the processing machinery, the large 24-*PHAS* precursors would accumulate in the cells where they were synthesized. This was indeed true as the *PHAS116* transcripts were found to be highly enriched in TAP at the 1.75 mm stage (this is a slightly later stage than in W23 anthers) in homozygous *dcl5 (dcl5-mu03//dcl5-mu03)* anthers (mixed inbred background) using smFISH (Fig. **2j**).

Overall, these data demonstrate that maize tapetal cells are the main site for 24-*PHAS* precursor accumulation and that accumulation starts at early meiotic prophase I and peaks during zygotene. We also established that accumulation is already substantial at the initial stages of meiosis in 1.25 mm anthers, approximately one day earlier than detected previously by analysis of 1.0 mm and 1.5 mm W23 whole anthers (Zhai *et al*., 2015).

### The spatial distribution of 24-nt phasiRNA biogenesis pathway components

Argonaute (AGO) proteins, binding partners of miRNAs and phasiRNAs, play essential roles in phasiRNAs biogenesis and RNA silencing (RNAi) (Czech & Hannon, 2011; Ma & Zhang, 2018). AGOs are assumed to be involved at two steps in 24-nt phasiRNA biogenesis and function: a specific AGO as the binding protein for the miR2275 trigger and one or several AGOs as the binding proteins for the diverse 24-nt phasiRNAs (Fig. **2a**). We identified transcripts encoding 18 AGO proteins in LCM cell collections (Fig. **3a**). Nine AGO protein transcripts were expressed in one or two cell collections, and the other nine AGO proteins were detected in all three cell collections. None of the AGO isotype transcripts were specific to tapetal cells. Fifteen of 18 AGO members were expressed in TAP at the 1.5 mm stage, providing a list of AGOs that could be involved in 24-nt phasiRNA biogenesis (Fig. **3a**). Among these, AGO1 family members have already been designated as candidates in previous studies in maize and rice (Fei *et al*., 2016; Kakrana *et al*., 2018; Araki *et al*., 2020), and now three AGO members (AGO10b, AGO2b, and AGO5c) not expressed in 1.5 mm TAP can likely be excluded as candidates for 24-nt phasiRNA biogenesis (Fig. **3a**). In meiocytes, transcripts were found for 11 AGO proteins, many of which were more highly expressed in ME than other cell types, but none were exclusive to ME (Fig. **3a**). Several of these AGO types were predicted in previous transcriptome profiling and spatial localization studies to be potential 24-nt phasiRNA binding partners, such as AGO18 family members (Zhai *et al*., 2015; Fei *et al*., 2016; Sun *et al*., 2019) and AGO2b (Fei *et al*., 2016) based on the timing of their expression. It is noteworthy that we detected abundant *AGO5c* transcripts, and these were enriched in ME and absent in TAP (Fig. **3a**). AGO5c is encoded by the maize *male-sterile28* gene and is homologous to rice MEL1 (Li *et al*., 2021), which is the binding partner for a subset of 21-nt phasiRNAs in rice (Komiya *et al*., 2014).

**Fig. 3.**
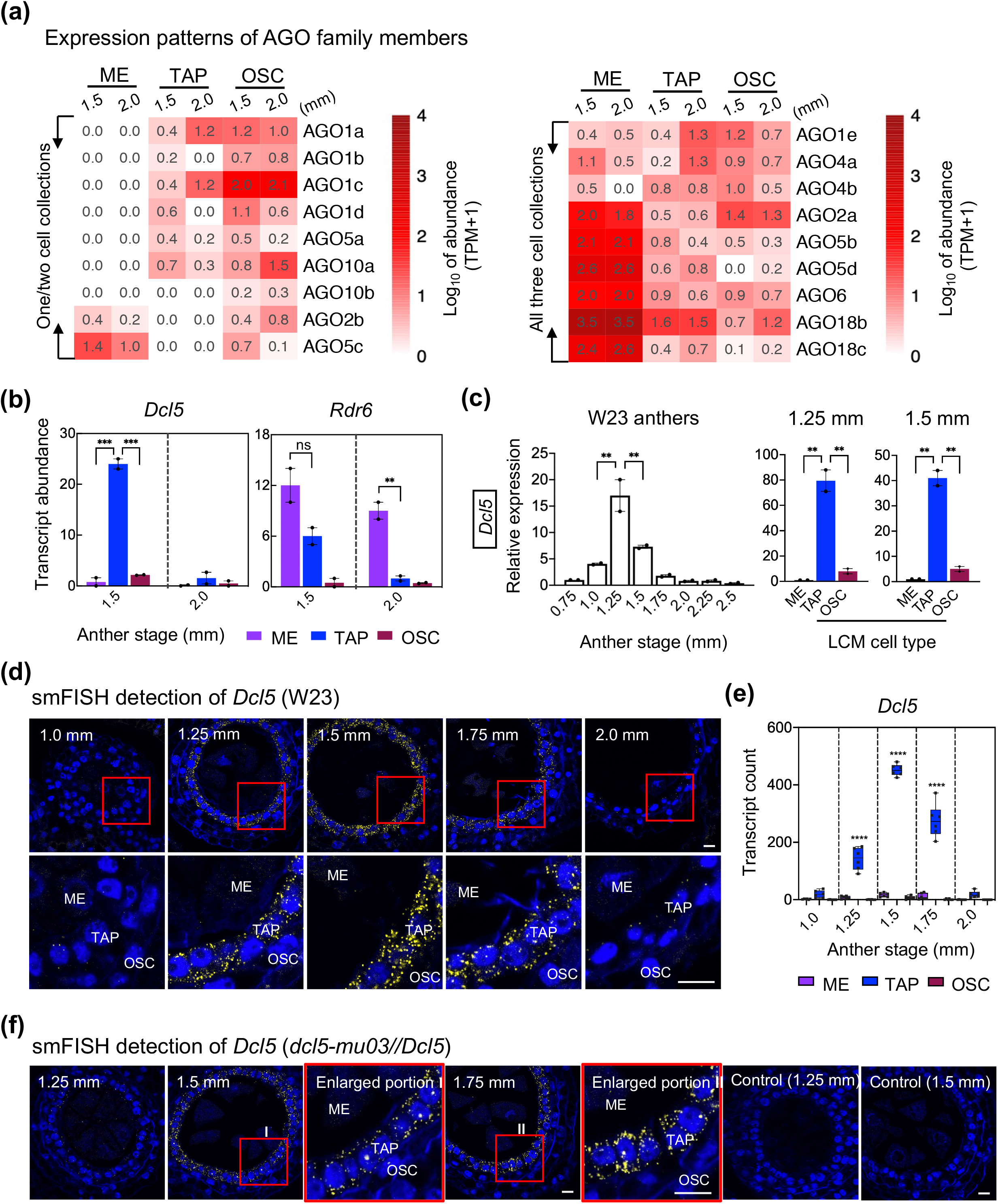
Evaluation of Argonaute, *Dcl5*, and *Rdr6* expression and *Dcl5* transcript localization in maize anthers. (**a**) RNA abundance from the 18 maize loci encoding Argonaute (AGO) proteins. At least one AGO participates in phasiRNA biogenesis by binding miR2275 and a few others have been shown to bind 24-nt phasiRNAs (see text). (**b**) Transcript abundances in LCM cells for two biogenesis genes from RNA-seq datasets. *Dcl5* was highly expressed in 1.5 mm TAP, and *Rdr6* was enriched in both ME and TAP. Differences among the LCM cell samples were considered significant for *P* values ≤ 0.05 (** *P* ≤ 0.01; *** *P* ≤ 0.001) using one-way ANOVA followed by Tukey tests. (**c**) RT-qPCR analysis for *Dcl5* at multiple anther stages and in LCM cell samples. The tapetal cell samples are significantly higher than ME and OSC (** *P* ≤ 0.01). Note that each graph has a different scale. (**d**-**e**) smFISH detection and quantification analysis for *Dcl5* transcripts in W23 maize anthers (1.0-2.0 mm). Probes are conjugated with Texas Red dye. The images below each panel are the enlarged portions of red rectangle areas. Yellow smFISH spots correspond to *Dcl5* transcripts are significantly higher in TAP (*** *P* ≤ 0.001). Scale bars = 10 μm. (**f**) smFISH detection of *Dcl5* transcripts in homozygous *dcl5-mu03//dcl5-mu03* anthers. Probes are conjugated with Texas Red dye. The red boxes designated as I and II indicate the area enlarged in the adjacent image. To validate the lack of *Dcl5* transcripts, no signal was detected in a control hybridization assay utilizing *Dcl5* probes applied to homozygous *dcl5-mu03//dcl5-mu03* anthers. Scale bars = 10 μm.

We investigated the transcript abundance of *Dcl5* and *Rdr6* in the LCM RNA-seq data. *Dcl5* was significantly higher in TAP at the 1.5 mm stage than any other samples (Fig. **3b**). In contrast, *Rdr6* was expressed in all three cell collections, with slightly higher expression levels in ME, followed by TAP at both stages (Fig. **3b**). To precisely pinpoint the timing of expression, RT-qPCR demonstrated that *Dcl5* transcripts were significantly higher in TAP samples (Fig. **3c**); furthermore, the expression pattern of *Dcl5* is similar to the three 24-*PHAS* precursors (*PHAS15, PHAS116*, and *PHAS296*), with the highest expression levels detected in TAP compared to ME or OSC at both the 1.25 mm and 1.5 mm stages (Fig. **3c**). As shown in Fig. **3d-f** with smFISH detection, *Dcl5* transcripts (yellow dots resulting from Texas Red labeled smFISH probes) were prominently localized in TAP at the 1.25 mm, 1.5 mm, and 1.75 mm stages in either W23 anthers or heterozygous *Dcl5*//*dcl5* fertile anthers. Far fewer yellow dots were detected in ME or OSC, and almost no signal was detected in the negative control samples (Fig. **3f**, Fig. **S6**). An additional smFISH analysis of *Dcl5* using AF647 labeled smFISH probes confirmed that *Dcl5* transcripts were localized mainly in TAP at 1.25 mm and 1.5 mm (Fig. **S7**). These data suggest that TAP is the main site for *Dcl5* accumulation at early meiotic prophase I in maize.

Previous studies have reported that four basic helix-loop-helix (bHLH) factors (*Ms23, Ms32, bHLH52*, and *bHLH122)* are required for normal tapetal cell specification and differentiation in maize (Moon *et al*., 2013; Nan *et al*., 2017). *Ms23* is the master transcription factor (TF) and is required for expression of 24-*PHAS* precursors, miR2275, and 24-nt phasiRNAs (Zhai *et al*., 2015). Here, *Ms23* and *bHLH51* were highly expressed in TAP at 2.0 mm, whereas *Ms32* and *bHLH122* were more abundant in TAP at 1.5 mm (Fig. **S8**). These TF expression patterns are consistent with our previous work in maize anthers, particularly the absence of *bHLH122* transcripts at 2.0 mm (Nan *et al*., 2017). Importantly, the TAP is significantly higher for these four TFs compared with the other two cell collections (Fig. **S8**). Collectively, the distribution of key TFs by RNA-seq and localization data provide further evidence that TAP is the main site for transcription of *PHAS* precursors and their processing into 24-nt phasiRNAs. Based on the timing of 24-*PHAS* accumulation, heterodimers of *Ms32* and *bHLH122* are candidates for direct regulation of 24-*PHAS* loci. Because *Ms23* is essential for expression of *bHLH122*, it is also a critical factor.

### 24-nt phasiRNAs are abundant in all three cell collections

To investigate the spatial distribution of 24-nt phasiRNAs in meiotic stage anthers, we collected ME, TAP, and OSC by LCM from 1.5 mm and 2.0 mm W23 maize anthers for low input sRNA-seq (Table **S4**, Table **S5**). We first classified sRNAs by length (18 nt to 30 nt) in FA and LCM cell collections. Confirming previous results (Zhai *et al*., 2015), the 24-nt size sRNA class is the most abundant at both stages (Fig. **S9a**). The 21-nt size sRNA class is also detectable, but with lower expression levels (Fig. **S9a**). Using the set of 118 24-*PHAS* loci for analysis (Fig. **2b**), 24-nt phasiRNAs from these loci were more abundant in all three cell collections at 2.0 mm than at the 1.5 mm stage (Fig. **4a**; Table **S6**). The expression patterns of 24-nt phasiRNAs and 22-nt miR2275 family members are very similar, with the highest abundance in ME at 2.0 mm during pachytene, followed by TAP at this stage (Fig. **4a**). A previous sRNA-FISH co-localization study reported that miR2275 and one 24-nt phasiRNA had higher abundances in AR/ME and TAP in maize anthers (Huang *et al*., 2019). Here, two of the miR2275 family members (miR2275a and miR2275b) were highly expressed, indicating they might play key roles in 24-*PHAS* precursor cleavage (Fig. **4b**). Although the precursors are most abundant in TAP cells (Fig. **2b, c**, Fig. **S2**), the 24-nt phasiRNAs accumulate to higher levels in the ME (Fig. **4a, c**, Fig. **S9b**). Individual 24-nt sRNA abundances from products of the same locus (*PHAS15, PHAS116*, and *PHAS296)* are not uniform: only a few of the 24-nt sRNAs show high abundance despite the expectation of equal stoichiometry from a single precursor molecule (Fig. **4d, e**, Fig. **S10a**). Furthermore, sRNAs localization studies illustrated that 24-nt phasiRNAs from three 24-*PHAS* loci (*PHAS15, PHAS116*, and *PHAS296*) accumulated in all three cell types, with higher expression levels in ME and TAP, lower in OSC from 1.25 mm to 2.0 mm in W23 anthers (Fig. **4f-h**, Fig. **S10b**). Almost zero expression, considered to be background, was detected in control hybridization assays (*dcl5* mutants, mouse HKH RNA control, and no labeled control) (Fig. **4h**, Fig. **S10c, d**). These distribution patterns lead us to hypothesize that the TAP is the site for nearly all 24-nt phasiRNAs biogenesis and that the presence of 24-nt phasiRNAs in ME and OSC depends on transport during early meiotic prophase I in maize.

**Fig. 4.**
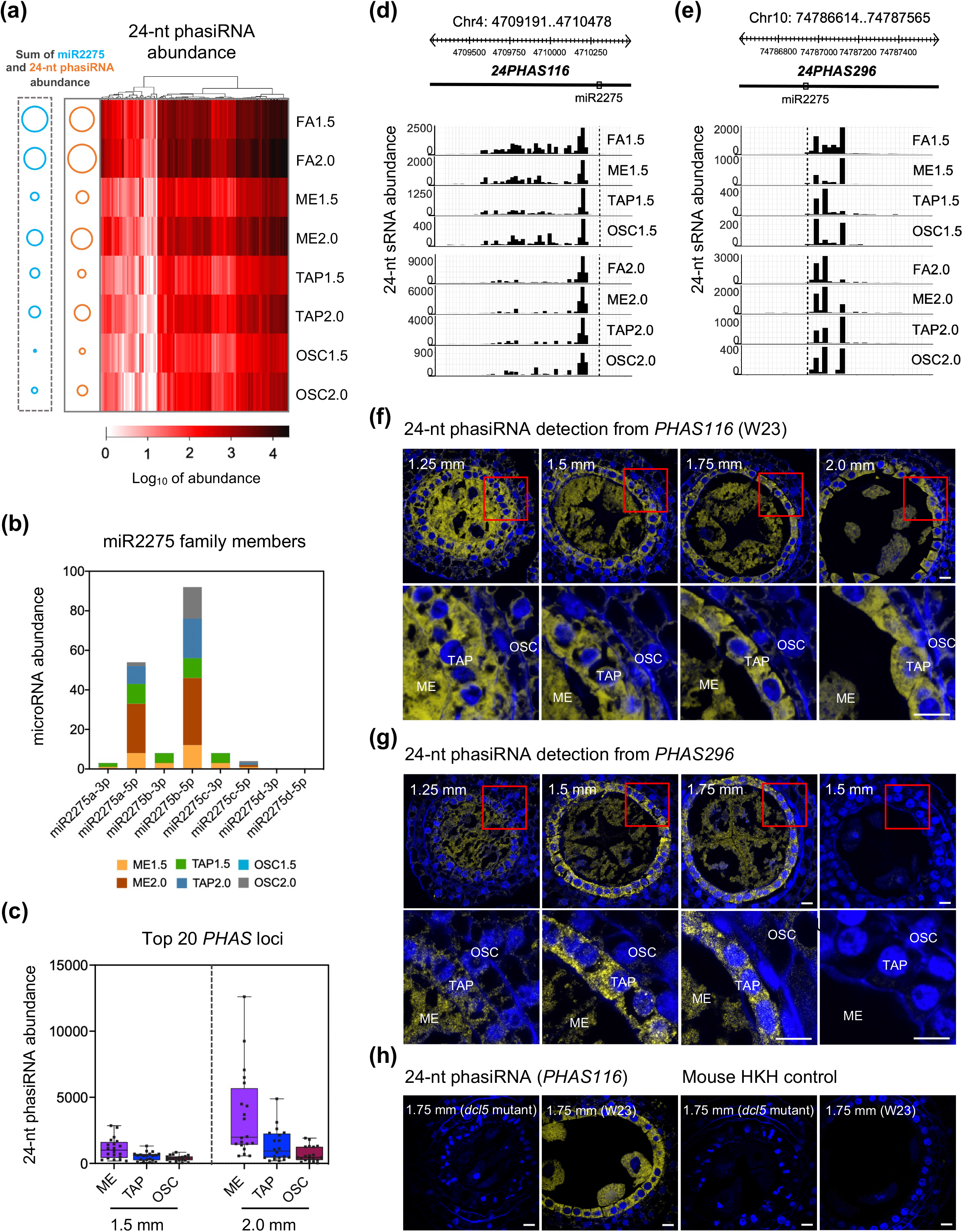
24-nt phasiRNA distribution in maize anthers. (**a**) Heat map depicting 24-nt phasiRNA abundance from 118 24-*PHAS* loci in fixed anthers and LCM cell sample collections. Solid bubbles on the left represent the total abundance of miR2275 family members and 24-nt phasiRNAs in that sample. The highest levels of 24-nt phasiRNAs and miR2275 family members are in 2.0 mm meiocytes, followed by tapetum at this stage. (**b**) The abundance of miR2275 family members in three LCM cell sample collections. (**c**) Box plot depicting the distribution of 24-nt phasiRNAs from the 20 most abundant 24-*PHAS* loci (same as panel **2c**). (**d**-**e**) 24-nt sRNA distributions from *PHAS116* and *PHAS296* in FA and LCM cells. The dotted lines represent the miR2275 cleavage sites, and each of the grid lines are spaced 24-nucleotide apart. The accumulation of 24-nt sRNA fragments generated from the same locus are not uniform: only a few of the 24-nt sRNAs show high abundance despite the expectation of equal stoichiometry from a single precursor molecule (Zhang *et al*., 2021). Overall, accumulation patterns are similar in fixed anthers and LCM cells at each locus. (**f**) 24-nt phasiRNA localization from *PHAS116* in W23 maize anthers (1.25-2.0 mm). Probes are conjugated with Texas Red dye. The red box in each image is enlarged in the panel immediately below each image. Yellow smFISH spots corresponding to 24-nt phasiRNAs are expressed in all three cell types. Scale bars = 10 μm. (**g**) 24-nt phasiRNA localization from *PHAS296* in W23 maize anthers (1.25-1.75 mm). Probes are conjugated with Texas Red dye. The red box in each image is enlarged in the panel immediately below each image. Yellow smFISH spots corresponding to 24-nt phasiRNAs are expressed in all three cell types. No signal was detected in an unlabeled control hybridization. Scale bars = 10 μm. (**h**) 24-nt phasiRNA localization from *PHAS116* in homozygous *dcl5-mu03//dcl5-mu03* anthers and W23 maize anthers at the 1.75 mm stage. Probes are conjugated with Texas Red dye. Low to zero signal was detected in negative control images with probes against mouse Hexokinase H (HKH) RNA (negative control). Scale bars = 10 μm.

## Discussion

In anthers, TAP is the innermost of the four somatic layers, adjacent to the germinal cells. The TAP plays essential roles in providing nutrients, enzymes, and pollen wall components to support AR, ME, and microspore (MSP) development. Defects in tapetal function or the abnormal differentiation of TAP usually leads to meiotic arrest or to later pollen abortion (Li *et al*., 2006; Xu *et al*., 2010; Moon *et al*., 2013; Niu *et al*., 2013; Nan *et al*., 2017; Teng *et al*., 2020). 24-nt phasiRNAs have been reported as abundant in meiotic stage maize anthers from hundreds of unique copy loci (Zhai *et al*., 2015). In TAP-defective, male-sterile mutants lacking key tapetal-expressed TFs, almost all the 24-nt phasiRNAs are missing (Zhai *et al*., 2015; Nan *et al*., 2017), which indicates that TAP is likely to be involved in 24-nt phasiRNA biogenesis. Our low-input RNA-seq data for ME, TAP, and OSC demonstrated that the TAP is the main source for the 118 24-*PHAS* precursor transcripts as well as *Dcl5* transcripts in 1.5 mm stage anthers (Fig. **2b, c**, Fig. **3b**), confirming a previous study in the same W23 background that 1.5 mm anthers were the peak stage for 24-*PHAS* loci production with 1.0 mm anthers lacking expression (Zhai *et al*., 2015). We further refined the timing using smFISH and RT-qPCR assays, demonstrating that three selected 24-*PHAS* loci (*PHAS15, PHAS116*, and *PHAS296)* and *Dcl5* transcripts were initially expressed at 1.25 mm during the leptotene-zygotene transition and were continuously expressed at 1.5 mm or 1.75 mm during zygotene (Fig. **2d-i**, Fig. **3d-f**, Fig. **S3**, Fig. **S7**). Moreover, the earlier timing for 24-nt phasiRNA generation corresponds to the stage just prior to and including the leptotene-zygotene transition (Fig. **4f, g**, Fig. **S10b**), which is a key checkpoint in meiosis. A previous study has reported that CHH DNA methylation is elevated in isolated ME, specifically at the phasiRNA-producing loci (Dukowic-Schulze *et al*., 2016). Therefore, a logical hypothesis is that some of the 24-nt phasiRNAs generated in the TAP and transported to ME might induce methylation in the corresponding loci. It is unclear how such *cis* methylation could contribute to meiotic regulation. Furthermore, in *dcl5* mutants, meiosis is normal in the near complete absence of 24-nt phasiRNAs (Teng *et al*., 2020), indicating that whatever role the 24-nt phasiRNAs play in meiosis, it is dispensable.

By using LCM-purified cell collections, our data demonstrate that TAP cells are the primary site of 24-*PHAS* precursor accumulation and 24-nt phasiRNA biogenesis. sRNA-seq data for ME, TAP, and OSC illustrated that 24-nt phasiRNAs and miR2275 family members were found in all three cell collections, with higher accumulation levels in ME at 2.0 mm during pachytene, followed by TAP at the same stage (Fig. **4a-c**). Furthermore, sRNA-FISH detection confirmed that the 24-nt phasiRNAs originating from three 24-*PHAS* loci (*PHAS15, PHAS116*, and *PHAS296)* were present in all three cell layers from 1.25 mm to 2.0 mm (Fig. **4f, g**, Fig. **S10b**). Our major new conclusion is the proposal that 24-nt phasiRNAs generated in the TAP move to other cell types. This conclusion is summarized in a model of 24-phasiRNA biogenesis and mobility in maize anthers during meiotic prophase I (Fig. **5**). We demonstrated that TAP is the main site of 24-nt phasiRNAs biogenesis, as TAP contain the vast majority of 24-*PHAS* precursors, *Dcl5* transcripts, adequate miR2275, and *Rdr6*. To understand why ME later contain the highest levels of 24-nt phasiRNAs, we note that there are approximately 30 tapetal cells touching each ME, whereas only approximately 0.6 tapetal cell touch neighboring OSC (Kelliher & Walbot, 2011). Utilizing the LCM cell numbers we recovered and the total 24-nt phasiRNA reads we generated (Table **S4**, Table **S6**), we calculated that each tapetal cell “donates” approximately 5% of its 24-nt phasiRNAs to ME or OSC during meiotic prophase I in maize anthers. If each TAP cells exports 5% (5 units) of its 24-nt phasiRNAs to ME cells, each ME would have 5 units x 30 = 150 units of 24-nt phasiRNAs, therefore reaching a higher level than the source TAP cells. In contrast, export of 5 units of 24-nt phasiRNAs to the OSC, would result in a low level of 24-nt phasiRNAs in the immediate receiving middle layer cell with further dilution as 24-nt phasiRNAs move throughout the OSC group. A challenge for future research will be documenting the path of 24-nt phasiRNA movement. Other cases of plant intercellular movement of sRNAs are probably through PD, regulating tissue or organ development by gene silencing and RdDM (Dunoyer *et al*., 2013). PD are membrane-lined intercellular channels that connect the cytoplasms of adjacent cells, such as between TAP and PMC/ME or non-TAP somatic cells in anthers (Mamun *et al*., 2005; Sager & Lee, 2014). Therefore, we speculate that 24-nt phasiRNAs move from the biogenesis site in TAP to ME or OSC through PD in meiotic prophase I maize anthers.

**Fig. 5.**
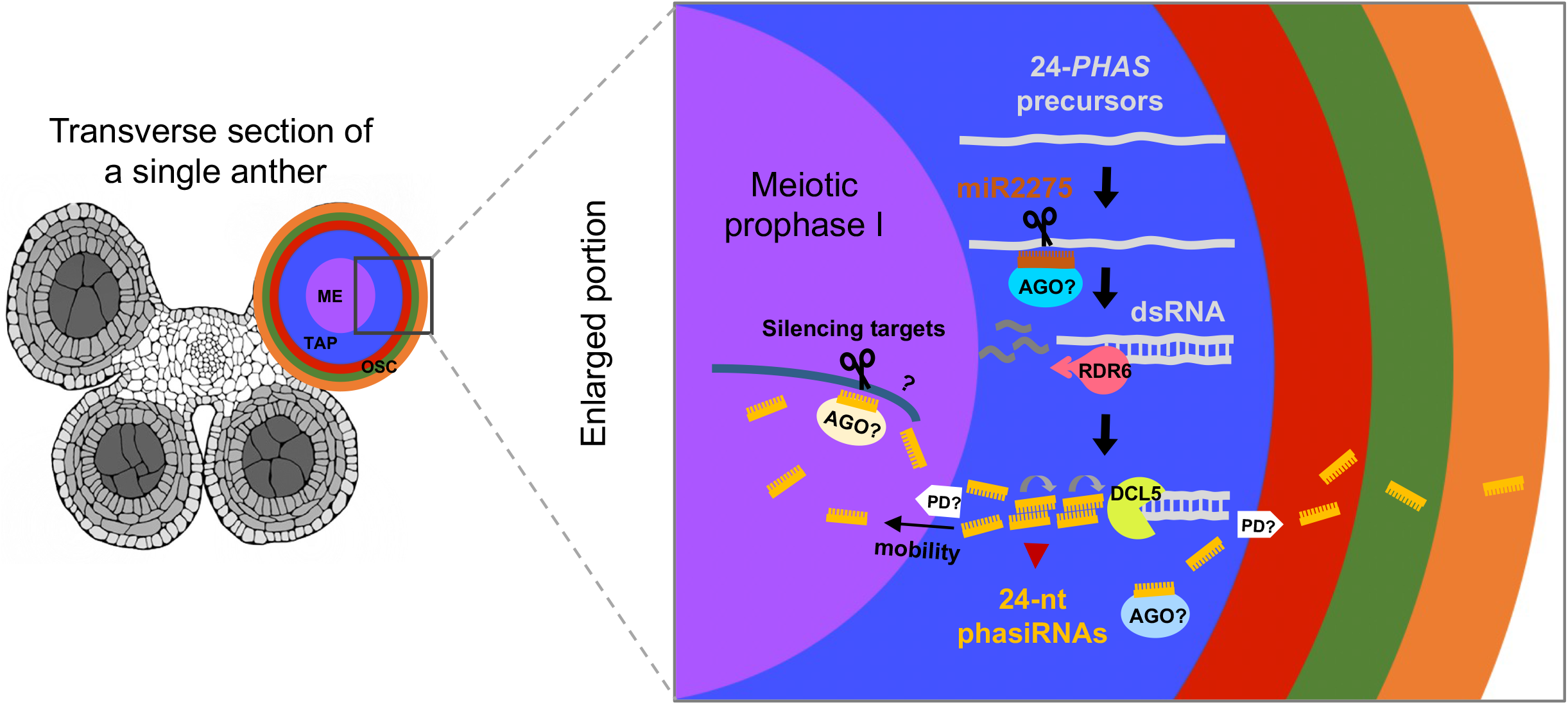
Model of 24-nt phasiRNAs biogenesis and mobility in maize anthers. On the left is a colorized confocal image as a transverse section of a single anther. On the right is an enlargement of a portion of a single lobe using the same color scheme. During the leptotene-zygotene transition, 24-nt phasiRNAs are generated in TAP and are proposed to move to other cell types. The 30 TAP cells in contact with each meiocyte are proposed to each export a small fraction of their 24-nt phasiRNAs to ME; similarly, a small amount of TAP 24-nt phasiRNA is exported to first to the adjacent Middle Layer (ML) and then distributed to other OSC.

In *Arabidopsis*, tapetal-derived 24-nt siRNAs induce DNA methylation in the germline and maintain genome integrity by silencing corresponding genes and transposons (Long *et al*., 2021). Here, 24-nt phasiRNAs are generated in TAP and then exported to ME (Fig. **5**). These exported 24-nt phasiRNAs are likely required for the elevated CHH DNA methylation at 24-*PHAS* loci in isolated ME in maize (Dukowic-Schulze *et al*., 2016). In *Arabidopsis*, the 24-nt siRNAs appear to have a broader role in modifying the ME genome and gene expression than the 24-nt phasiRNAs have in maize. This is particularly true for any impact 24-nt phasiRNAs might have on gene expression, because no mRNA targets for this class of small RNAs have been identified. Nonetheless, it is interesting that small RNAs synthesized in the TAP are transferred to maize ME. Analysis of *dcl5* maize mutants establishes that in the absence of 24-nt phasiRNA biogenesis, tapetal cells fail to differentiate properly at normal maize growing temperatures (28 °C), while meiosis proceeds normally (Teng *et al*., 2020). These observations indicate that the 24-nt phasiRNAs play an essential role in the TAP -- not in ME -- during the meiotic stages; later, microspore failure renders *dcl5* mutants male-sterile, a fact that we attribute to the failure of the tapetum to supply nutrients and wall components to the developing uninucleate microspores. Mutants grown at low temperature (21 °C) are fertile and tapetal development is slow but sufficient indicating that 24-nt phasiRNAs are not required under these conditions. Considering these observations *in toto*, it is possible that in maize the transfer of 24-nt phasiRNAs to meiocytes is an evolutionary relic that no longer serves (or fully serves) a required function and that the primary role of the 24-nt phasiRNAs is to support rapid tapetal differentiation.

## Acknowledgements

We thank M. Zhang for initial suggestions on project design and for instructing X. Zhou in the LCM technique; B. Nelms for help with low input RNA-seq library construction and raw data processing; Y. Chan and Y. Liu for help in data handling; J. Caplan for helpful advice on smFISH; J. Zhan assisted retrieval and organization of the sequences of 24-*PHAS* precursors and 24-nt phasiRNAs; and J. Dinneny for use of the SP8 confocal microscope. This research was supported by National Science Foundation award 17540974. Requests for materials should be addressed to.

## Author Contributions

X.Z., B.C.M., and V.W. conceived and designed the project. X.Z. performed most of the experiments and bioinformatics. K.H. and M.B. designed and synthesized smFISH probes. K.H., M.B., A.A., T.K., and T.C. performed smFISH and imaging analysis. T.C. constructed sRNA libraries and performed raw data processing. X.Z. and V.W. wrote the manuscript with editing by B.C.M. and K.H. and with contributions from all co-authors.

## Data Availability

Sequencing data were deposited to the Gene Expression Omnibus (accession no. GSE182588 for RNA-seq data; accession no. GSE182476 for sRNA-seq data) and will be made public after peer review.

## Competing interests

The authors declare no competing interests.

## Supporting Information

**Fig. S1** Sequential application of LCM to isolate ME, TAP, and OSC in 1.5 mm and 2.0 mm W23 anthers.

**Fig. S2** Transcript abundances of three 24-*PHAS* loci (*PHAS15, PHAS116*, and *PHAS296)* in LCM cell sample collections in RNA-seq data.

**Fig. S3** *PHAS15* localization in maize anthers.

**Fig. S4** Negative control hybridization assay.

**Fig. S5** *ZmCyanase* localization in maize anthers.

**Fig. S6** No labeled control hybridization assay.

**Fig. S7** *Dcl5* localization in maize anthers.

**Fig. S8** The distribution of four tapetal-enriched TFs (*Ms23, Ms32, bHLH51*, and *bHLH122)* in LCM cell sample collections at 1.5 mm and 2.0 mm in the RNA-seq dataset.

**Fig. S9** sRNA distribution in maize anthers.

**Fig. S10** 24-nt phasiRNAs localization in maize anthers.

**Table S1** Primer sequences for CEL-seq 2 library preparation.

**Table S2** Primer sequences for RT-qPCR by the TaqMan assay.

**Table S3** Probes sequences (5’-> 3’) used for smFISH or sRNA-FISH.

**Table S4** List of LCM cell sample collections from 1.5 mm and 2.0 mm maize anthers.

**Table S5** Summary of RNA-seq and sRNA-seq libraries prepared from FA and LCM cell sample collections.

**Table S6** Coordinates and abundance of the 126 24-*PHAS* loci and their corresponding 24-nt phasiRNAs in FA and LCM cells.

## Notes

### Competing Interest Statement

The authors have declared no competing interest.

### Summary of Updates

Figure 4 revised. Supplemental files updated.

