## Supporting Information for "24-nt phasiRNAs move from tapetal to meiotic cells in maize anthers"

*New Phytologist* **Supporting Information**

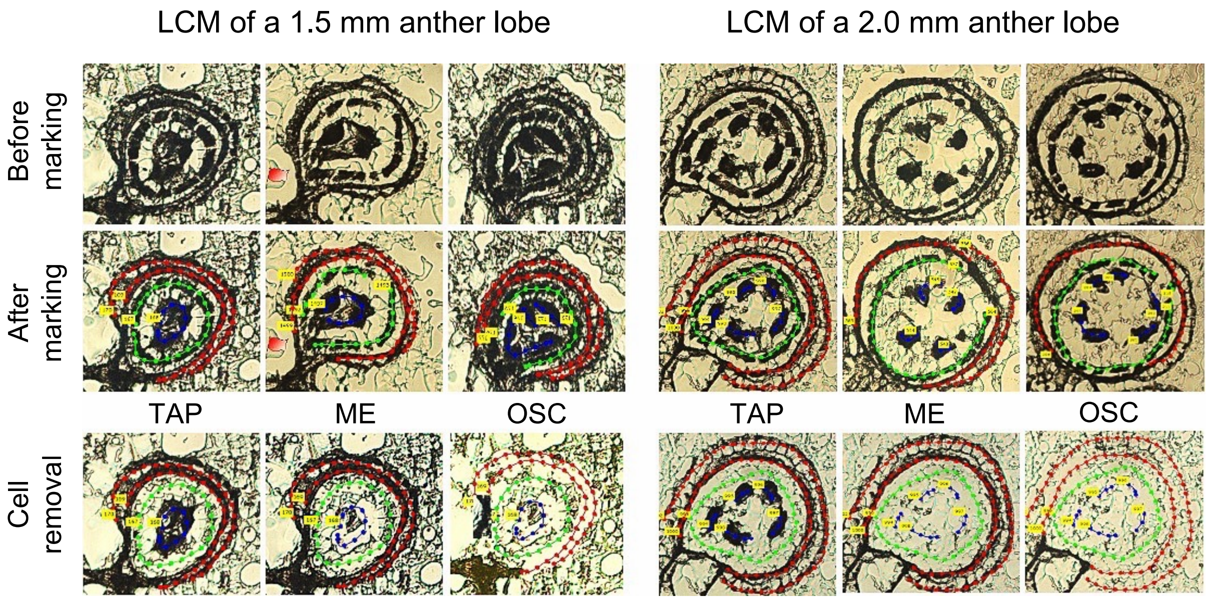


**Fig. S1 Sequential application of LCM to isolate ME, TAP, and OSC in 1.5 mm and 2.0 mm W23 anthers.**

ME marked in blue dotted lines, TAP marked in green dotted lines, and OSC marked in red dotted lines. At the end of each dotted line, the PALM software autogenerates a yellow box that tags the cell-layer with a number. By choosing the appropriate number, three different cell layers were collected separately into PCR tubes with RNA extraction buffer.

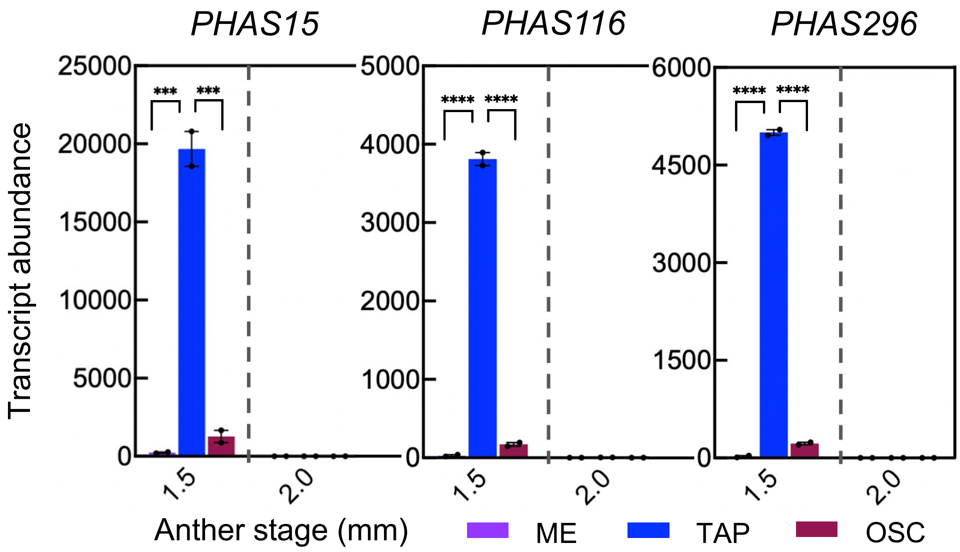


**Fig. S2 Transcript abundances of three *24-PHAS* loci (*PHAS15*, *PHAS116*, and *PHAS296*) in LCM cell sample collections in RNA-seq data.**

The tapetal cell samples are significantly higher than ME and OSC in each group. Prism 8 software was applied to determine *P* values with one-way ANOVA and Tukey tests. Differences among the samples were considered significant for *P* values $\leq 0.05$ (* $P \leq 0.05$; ** $P \leq 0.01$; *** $P \leq 0.001$; **** $P \leq 0.0001$). Note that there is a different scale for each 24-*PHAS* locus.

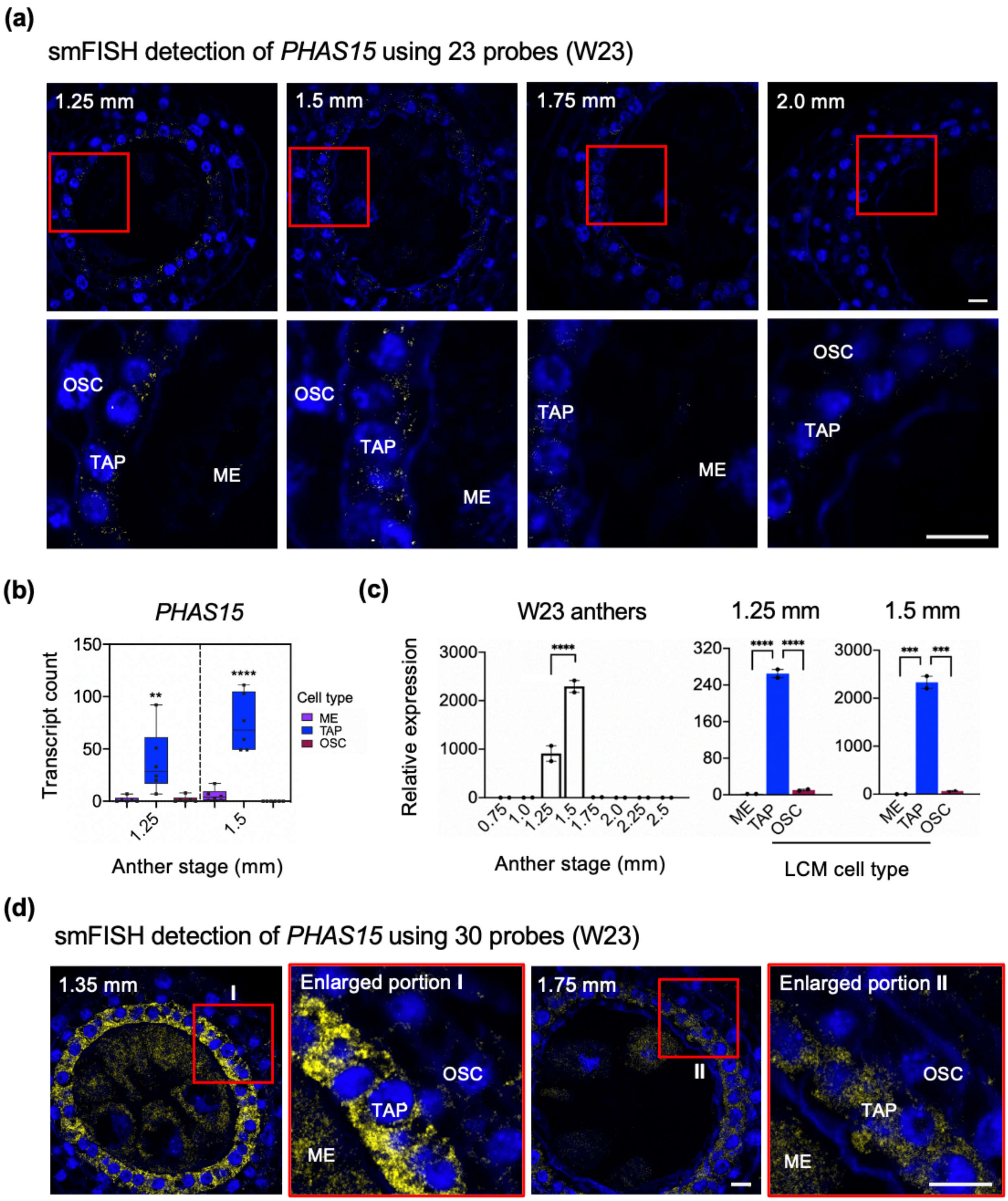
(a)
smFISH detection of PHAS15 using 23 probes (W23)
1.25 mm
1.5 mm
1.75 mm
2.0 mm
OSC
TAP
ME
(b)
PHAS15
Transcript count
Cell type
ME
TAP
OSC
Anther stage (mm)
(c)
W23 anthers
1.25 mm
1.5 mm
Relative expression
Anther stage (mm)
LCM cell type
(d)
smFISH detection of PHAS15 using 30 probes (W23)
1.35 mm
Enlarged portion I
1.75 mm
Enlarged portion II

**Fig. S3 *PHAS15* localization in maize anthers.**

(**a**) smFISH was performed using 23 probes (excluded were 7 probes that overlap with 24-nt phasiRNAs) to localize *PHAS15* in W23 maize anthers (1.25-2.0 mm). Probes are conjugated with dye AF647. The red box in each panel is enlarged in the image immediately below. Yellow smFISH spots correspond to *PHAS15* transcripts, which are highly enriched in TAP. Probes were designed with the negative strand sequence of *PHAS15.* Scale bars = 10 µm. (**b**) Quantification analysis of images in (**a**) at the 1.25 mm and 1.5 mm stages. The tapetal cells are significantly higher (** $P \leq 0.01$; **** $P \leq 0.0001$). (**c**) RT-qPCR results for *PHAS15* in whole anthers (0.75-2.5 mm) and LCM cells (1.25 and 1.5 mm). The tapetal cell samples are significantly higher (*** $P \leq 0.001$; **** $P \leq 0.0001$) than ME or OSC. Note that there is a different scale for this locus. (**d**) smFISH was performed using 30 probes (included 7 probes that overlap with 24-nt phasiRNAs) to localize *PHAS15* in W23 maize anthers (1.35 mm and 1.75 mm). Probes are conjugated with dye AF647. The enlarged portions in the red boxes are from the left images. Yellow smFISH spots correspond to *PHAS15* transcripts or the overlapped 24-nt phasiRNAs are detected in all cell types. Scale bars = 10 µm.

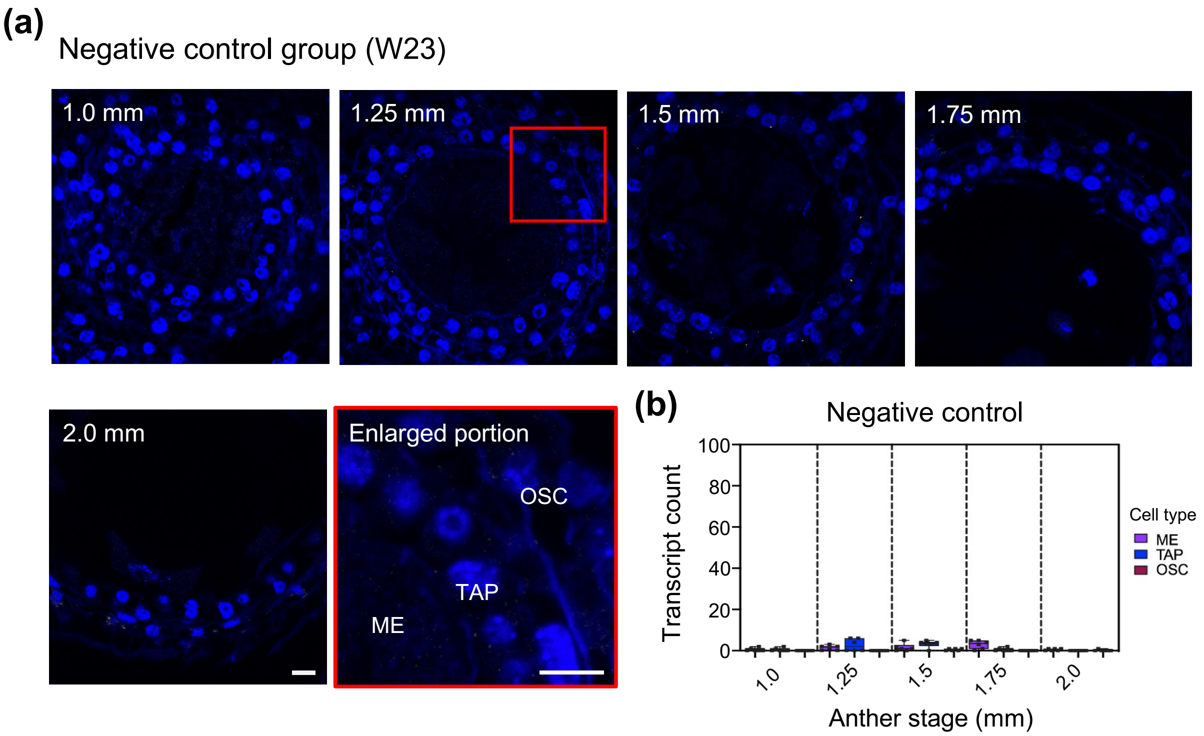


**Fig. S4 Negative control hybridization assay.**

(**a**) Negative smFISH images in W23 maize anthers (1.0-2.0 mm). Probes were designed with the positive strand sequence of *PHAS15* and conjugated with dye AF647. The red box in the 1.25 mm sample was enlarged and is shown immediately below the image. Scale bars = 10 µm. (**b**) Quantification analysis of images in (**a**).

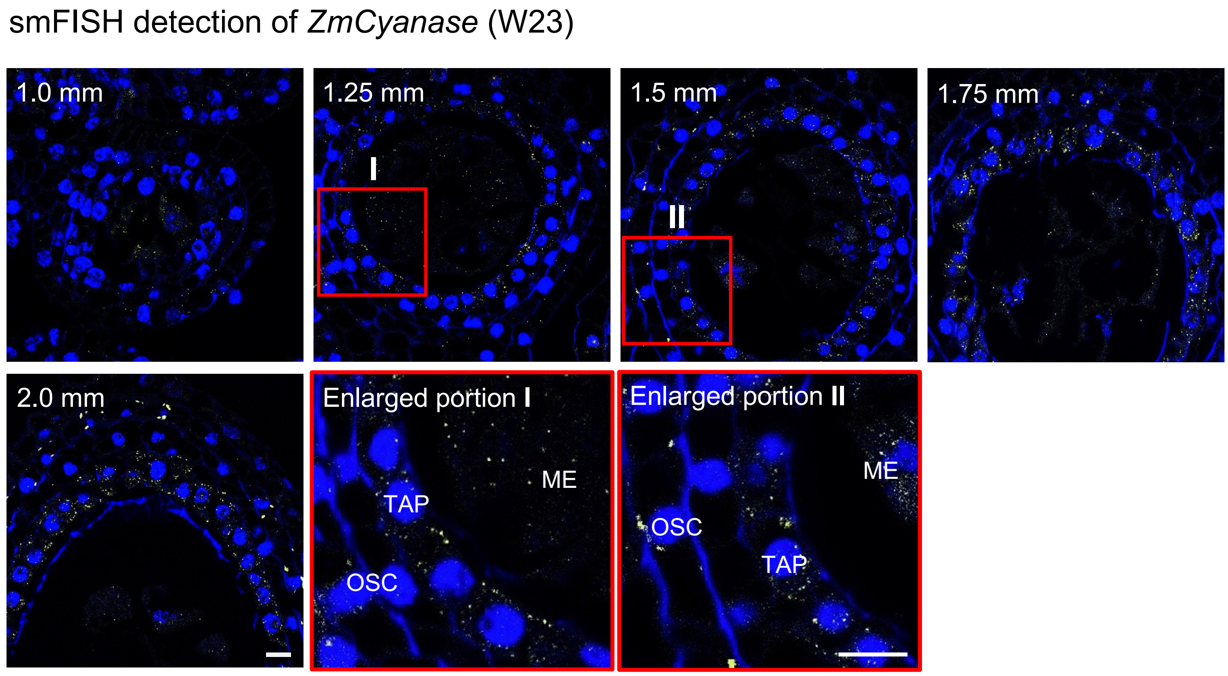


**Fig. S5 *ZmCyanase* localization in maize anthers. smFISH was performed to localize *ZmCyanase* in W23 maize anthers (1.0-2.0 mm).**

Probes were conjugated with Texas Red dye. The red boxes labeled I and II in the 1.25 mm and 1.5 mm samples were enlarged and are shown immediately below the corresponding image. Yellow smFISH spots corresponding to *ZmCyanase* transcripts were detected in all three cell types. Scale bars = 10 µm.

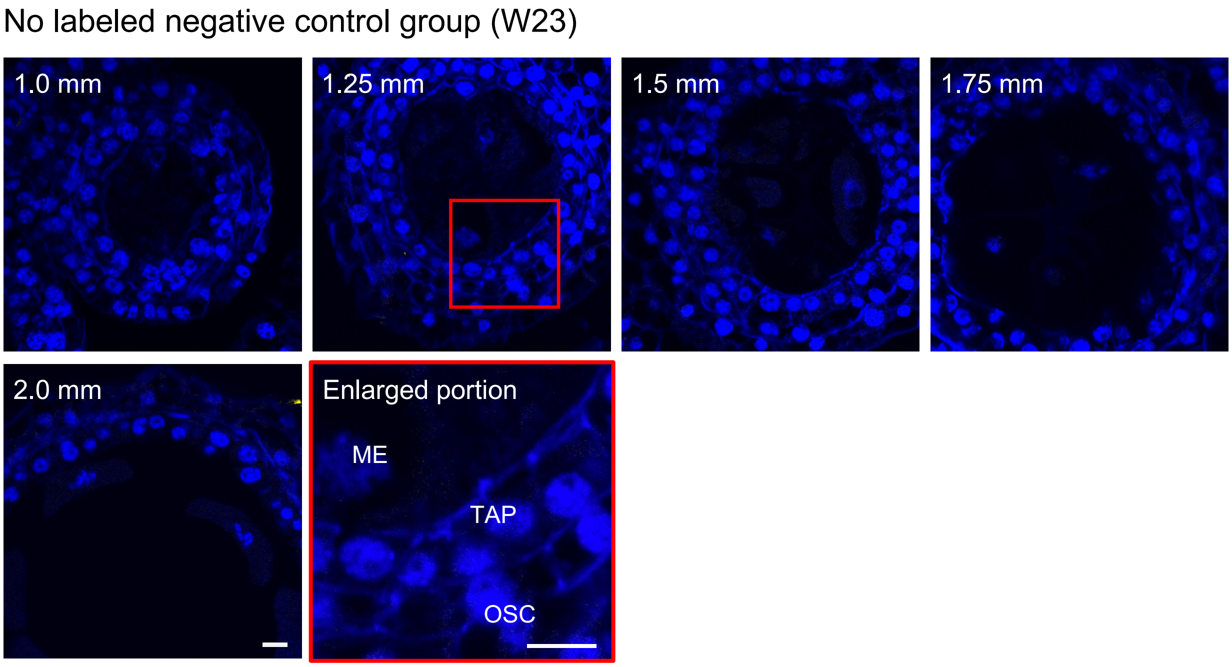


**Fig. S6 No labeled control hybridization assay**.

Negative control images with unlabeled smFISH probes in W23 maize anthers (1.0-2.0 mm). The red box in the 1.25 mm sample was enlarged and is shown immediately below the image. No signals were detected. Scale bars = 10 µm.

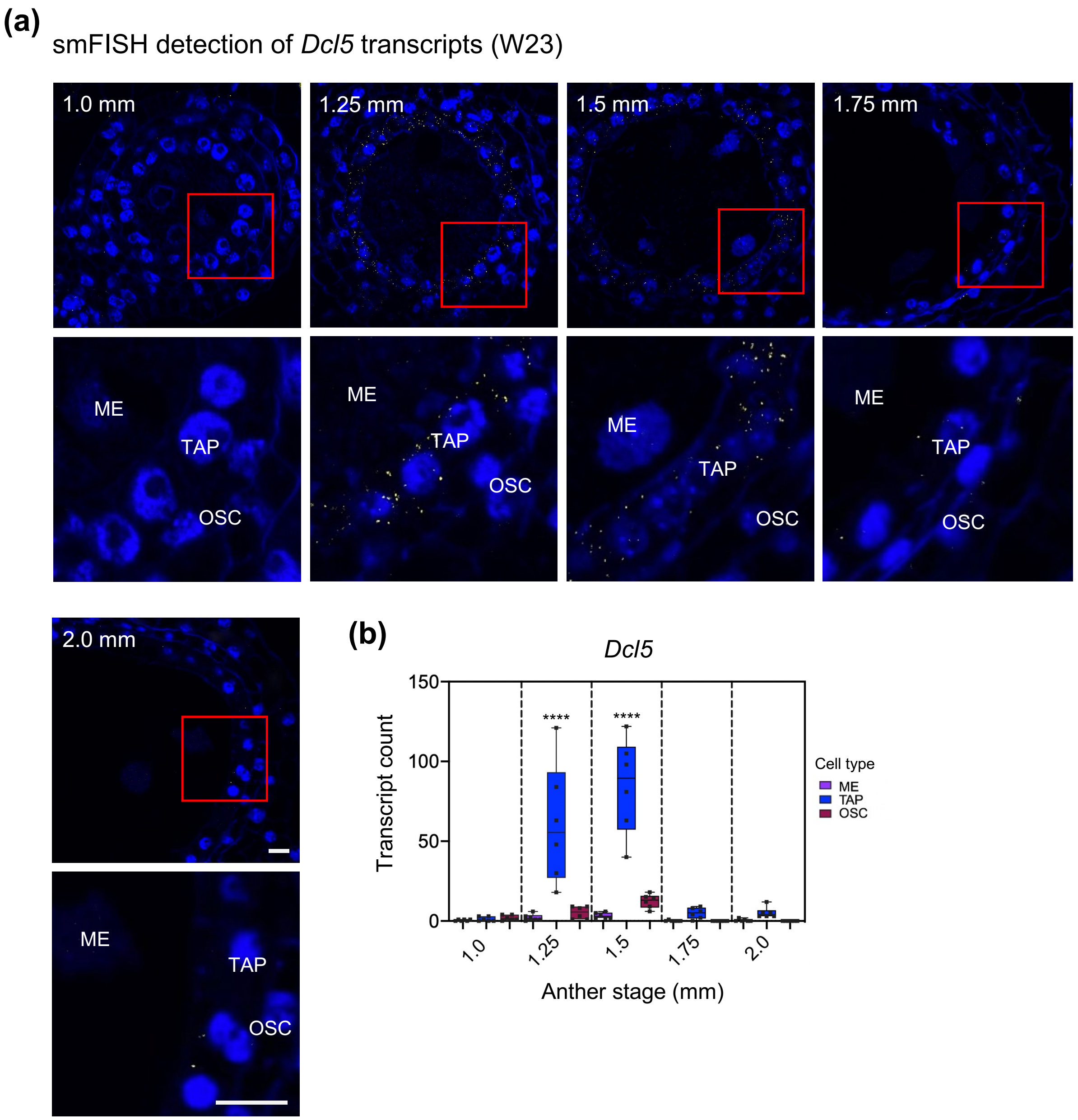


**Fig. S7 *Dcl5* localization in maize anthers.**

smFISH detection (**a**) and quantification analysis (**b**) for *Dcl5* transcripts in W23 maize anthers (1.0-2.0 mm). Probes are conjugated with dye AF647. The red boxes in each image were enlarged and are shown directly below the corresponding image. Yellow smFISH spots corresponding to *Dcl5* transcripts are significantly higher in TAP (**** $P \leq 0.0001$). Scale bars = 10 μm.

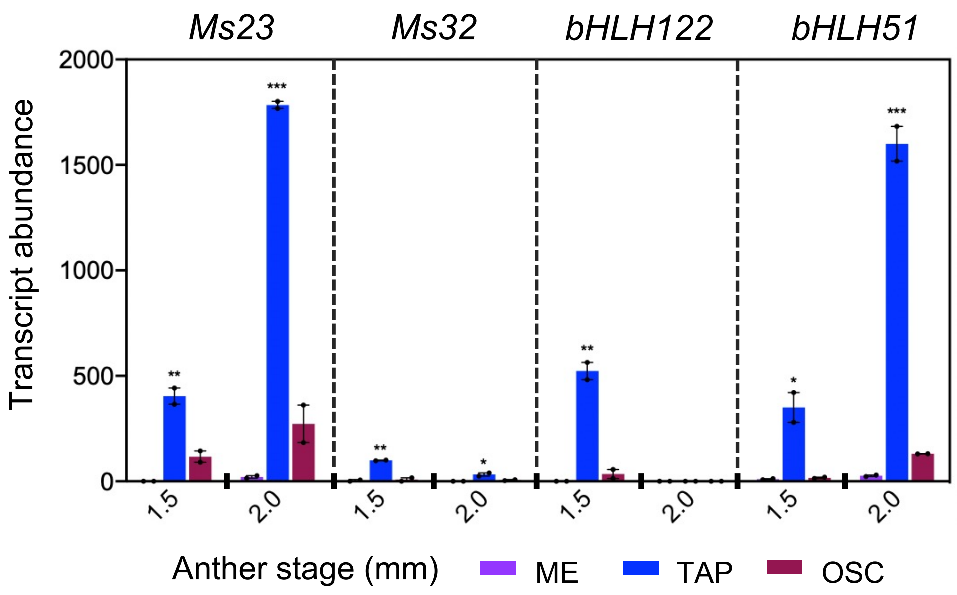


**Fig. S8 The distribution of four tapetal-enriched TFs (*Ms23*, *Ms32*, *bHLH51*, and *bHLH122*) in LCM cell sample collections at 1.5 mm and 2.0 mm in the RNA-seq dataset.**

The tapetal cell samples are significantly higher (* $P \leq 0.05$; ** $P \leq 0.01$; *** $P \leq 0.001$) than ME and OSC at each stage.

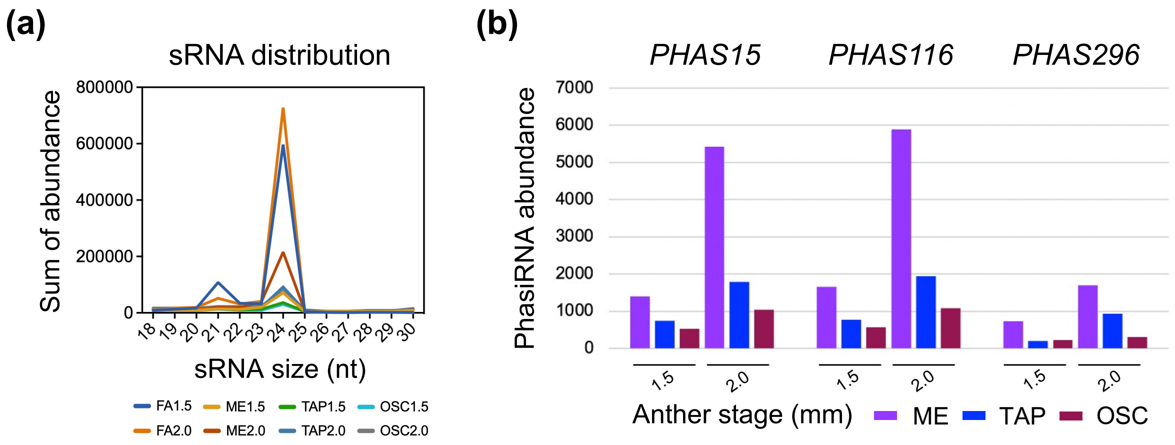


**Fig. S9 sRNA distribution in maize anthers.**

(**a**) Line plot showing the abundance distribution of different size classes (18-30 nt in length) of sRNAs in fixed anthers and LCM cell sample collections in 1.5 mm and 2.0 mm stage maize anthers. (**b**) The distribution of 24-nt phasiRNAs generated from three 24-*PHAS* precursors (*PHAS15*, *PHAS116*, and *PHAS296*) in 1.5 mm and 2.0 mm maize anthers.

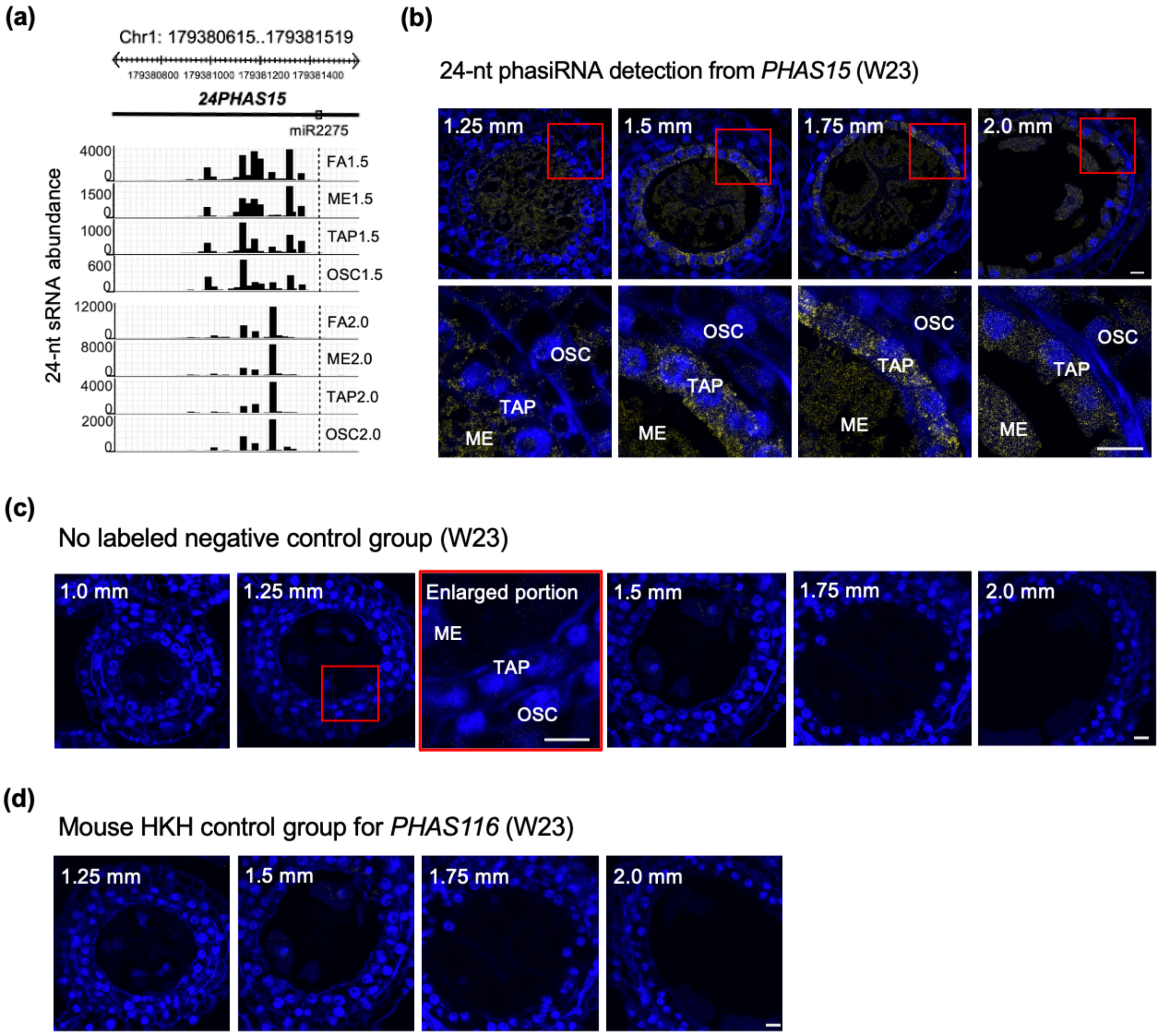


**Fig. S10 24-nt phasiRNAs localization in maize anthers.**

(**a**) 24-nt sRNA distributions from *PHAS15* in FA and LCM cell sample collections from 1.5 mm and 2.0 mm stage maize anthers. (**b**) 24-nt phasiRNAs localization from *PHAS15* in W23 maize anthers (1.25-1.75 mm). The red boxes in each image were enlarged, and these are shown directly below the corresponding image. Yellow smFISH spots corresponding to 24-nt phasiRNAs are expressed in all three cell types. Scale bars = 10 µm. (**c**) Negative control images with unlabeled sRNA-FISH probesin W23 maize anthers (1.0-2.0 mm). The red box in the 1.25 mm sample was enlarged and is shown to the right of this image. Scale bars = 10 µm. (**d**) Negative control images with probes against mouse Hexokinase H (HKH) RNA in W23 maize anthers (1.25-2.0 mm). Scale bars = 10 µm.
